# Theranostic gold in a gold cage nanoparticle for photothermal ablation and photoacoustic imaging of skin and oral infections

**DOI:** 10.1101/2023.05.05.539604

**Authors:** Maryam Hajfathalian, Christiaan R. de Vries, Jessica C. Hsu, Ahmad Amirshaghaghi, Yuxi C. Dong, Zhi Ren, Yuan Liu, Yue Huang, Yong Li, Simon Knight, Pallavi Jonnalagadda, Aimen Zlitni, Elizabeth Grice, Paul L. Bollyky, Hyun Koo, David. P. Cormode

## Abstract

Biofilms are structured communities of microbial cells embedded in a self-produced matrix of extracellular polymeric substances. Biofilms are associated with many health issues in humans, including chronic wound infections and tooth decay. Current antimicrobials are often incapable of disrupting the polymeric biofilm matrix and reaching the bacteria within. Alternative approaches are needed. Here, we describe a unique structure of dextran coated gold in a gold cage nanoparticle that enables photoacoustic and photothermal properties for biofilm detection and treatment. Activation of these nanoparticles with a near infrared laser can selectively detect and kill biofilm bacteria with precise spatial control and in a short timeframe. We observe a strong biocidal effect against both Streptococcus mutans and Staphylococcus aureus biofilms in mouse models of oral plaque and wound infections respectively. These effects were over 100 times greater than that seen with chlorhexidine, a conventional antimicrobial agent. Moreover, this approach did not adversely affect surrounding tissues. We conclude that photothermal ablation using theranostic nanoparticles is a rapid, precise, and non-toxic method to detect and treat biofilm-associated infections.

## Introduction

Biofilms, communities of bacteria within an extracellular polymeric matrix (EPS), are implicated in many chronic infections (1–3). Once biofilms are established, they are notoriously difficult to eradicate due to the presence of an extracellular matrix which surrounds bacteria and protects them from antibiotics (4, 5). Hence, it is important to locate the biofilms for rapid and precise action against them. Current therapeutic modalities struggle to disrupt biofilms and often fail to efficiently kill the microbes within (6, 7). Consequently, biofilm-associated infections are responsible for terrible human suffering and massive economic expense (8). The development of therapies capable of killing biofilm-associated bacteria would have a tremendous impact on the treatment of chronic infections.

Two settings where biofilms are relevant to human health are the skin and the mouth. Dental caries afflicts nearly half of the world’s population and is a major public health problem according to the US Centers for Disease Control (9). One of the major oral pathogens involved in dental caries is Streptococcus mutans (S. mutans); its ability to form biofilms is a key aspect of this pathology (10, 11). Over the past decade researchers have devised innovative antimicrobial strategies to inhibit S. mutans biofilm formation and disrupt extracellular polymeric substances (EPS), the building blocks of biofilms (12–14). Unfortunately, antibacterial resistance, a lack of appropriate diagnostic tools, and a paucity of convenient and affordable treatments remain barriers to treating oral biofilms (15, 16).

Wound infections affect 6.5 million people in the U.S., with estimated annual costs of $28 billion (17). Bacterial biofilms are a major factor in wound chronicity due to effects on delayed healing and impaired bacterial clearance by antibiotics (18, 19). In most contexts, Staphylococcus species are the most common bacteria recovered from chronic wounds. There are many strategies to clear wound biofilms with periodic wound debridement being the most common (20, 21). However, wound biofilms remain a major unsolved, growing problem (22).

Photothermal therapy (PTT) is an antibacterial strategy which utilizes light sources to activate nanoagents and convert the light energy into heat, killing the pathogens (23, 24). This antibacterial strategy is non-invasive and effective against biofilm bacteria, including those resistant to conventional antibiotic drugs (25–27). However, there is still a need to develop anti-biofilm agents that are photothermally efficient, biocompatible, and photostable (28–30). Advances in shape engineering, synthesis methods, and coatings of nanoparticles, and their combinations with organic materials have introduced novel functionalities for nanostructures and their utility in different bioapplications and therapeutic methods (31, 32). Gold nanoparticles in particular have been used to enhance antibiotic efficacy and reduce bacterial viability in vitro (33–35). Optically tuned gold nanoparticles with different sizes and structures have been also studied for their use in photothermal ablation (36, 37). Gold nanorods, nanostars and nanoshells have shown strong antibacterial effects against bacterial biofilms (38–40). In addition, gold nanocages with hollow interior and thin porous walls have been used as PTT agents (41–43). However, these nanoparticles need to be engineered to have a strong extinction spectrum in NIR region and higher photothermal efficiency to be capable of killing bacteria effectively and treating infections in a short time frame. Moreover, pre-clinical studies (specially for oral infections) are lacking for most of these agents. There is an urgent need for ex vivo and in vivo demonstrations of efficacy that would allow further development of PTT(14, 44).

There is also a need to visualize biofilm infections on wounds, teeth, and gums in order to define the infected lesions precisely, determine the depth, and document ablation in real-time (45–47). Photoacoustic imaging (PA) is a powerful imaging technique that allows detection of bacterial infection (48, 49). PA utilizes photothermal expansion of light absorbing contrast agents to generate ultrasound waves under pulsed laser irradiation. PA has less background, higher spatial resolution, stronger contrast, better deep tissue penetration than optical imaging (50, 51). The development of a multi-function platform that could image oral and wound bacterial infection as well as ablate bacterial infections simultaneously would be a boon to infection control efforts.

Here, we report multi-functional gold in a gold cage photothermal nanoparticles (PTNP), which specifically detect biofilm and kill bacteria with high spatial precision in less than a minute. These nanoparticles were formed by growth of a silver shell over a gold core and subsequent galvanic replacement of the silver with a gold cage. This morphology has a unique plasmon peak in NIR which provides them an excellent performance in biomedical applications involving optical excitation or transduction. We have evaluated the photothermal properties, biofilm uptake, antimicrobial action, cytotoxicity, and photoacoustic imaging ability of these nanoparticles. Our in vitro data demonstrate high potency to eradicate biofilms and eliminate wound and oral pathogens without toxicity to eukaryotic cells. Furthermore, PTNP was examined for its efficacy as an anti-biofilm agent to detect and treat in vivo biofilms in wound infections and on teeth using rodent models. These exciting findings demonstrate the potential of these novel nanostructures for therapeutic applications.

## Results

### Synthesis of gold in a gold cage photothermal nanoparticles

PTNP were synthesized through the following method: a modified Turkevich method was used to develop small seeds (52). A seeded growth method was then used to make larger nanoparticles with the equilibrium shape (a truncated octahedron enclosed by six square facets and eight hexagonal facets (53); which we term cores in this manuscript). We then performed a reduction of silver nitrate (AgNO_3_) onto the cores to produce ‘core in a shell’ structures followed by a galvanic replacement reaction (GRR) of silver with gold ions (HAuCl_4_) to change the shape of the shells into nanocages, which we termed ‘gold in a gold cage’. Figures 1A and B show a schematic depiction and transmission electron microscopy (TEM) image of the PTNP, respectively.

**Figure 1.**
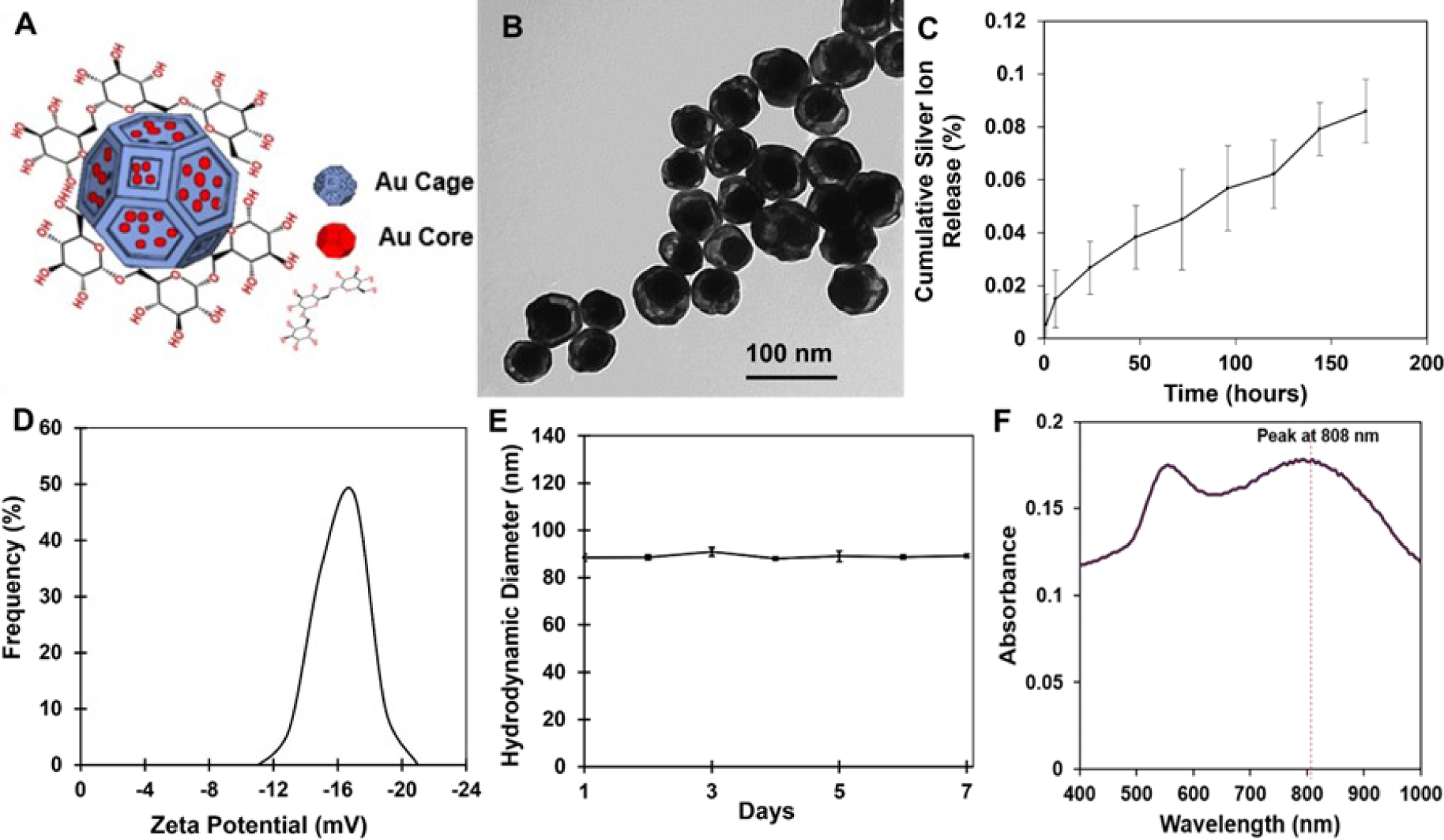
PTNP with unique morphologies are stable and alternative plasmonic nanoagents for biomedical applications. (**A**) Schematic depiction of PTNP structure, which consists of an Au core encapsulated by Au cage and coated with DEX. (**B**) TEM of PTNP (Scale bar: 100 nm). (**C**) Silver ion release from PTNP when incubated in DI water at 37 °C over a period of 7 days (n=6). (**D**) Zeta Potential data of DEX-coated PTNP. (**E**) Hydrodynamic diameter of PTNP in DI water during a week (n=4). (**F**) UV-vis spectrum of DEX-PTNP. The data were presented as mean ± SD.

For this experiment, the small gold nanoparticles (Au seeds,16 ± 2 nm) acted as nucleation sites and a seeded-growth synthesis method was used to synthesize the larger seeds (Au core, 78 ± 3 nm). Then, AgNO_3_ was reduced with ascorbic acid (AA) in a solution of Au cores to make a core@shell structure (Au@Ag, 86 ± 6 nm) (Figure S1). The thickness of these shells was measured using TEM and was found to be 8 ± 1 nm. The size of these is tunable based on the reaction time, the amount of reducing agent, and the concentration of AgNO_3_. The final step of the shape engineering to create gold in a gold cage structure was adding HAuCl_4_ to the Au@Ag structure solution to initiate GRR, which is a spontaneous oxidation–reduction reaction where HAuCl_4_ is reduced, and silver is oxidized. This reaction caused the formation of cages, which we termed Au cages, with a size of 85 ± 5 nm around the Au cores. The composition of the PTNP structures is 87 ± 2% Au and 13 ± 4% Ag, as measured by inductively coupled plasma-optical emission spectroscopy (ICP-OES) analysis. These results were also confirmed with energy dispersive X-ray spectroscopy (EDS) measurements (Figure S2A). The low amount of silver on cages as well as its being alloyed with Au can be expected to suppress possible silver ion release, improve the overall stability, and extend the lifetime of the anti-biofilm agent. Figure 1C shows that silver ion release from PTNP over 7 days is very low (less than 0.085%), and it is much lower than that found for pure silver nanoparticles (54). This result agrees with prior work that showed that alloying silver with gold suppressed silver ion release (55).

Finally, a surface coating was added to stabilize the PTNP structures and enhance their biocompatibility and uptake by biofilms (56, 57). The DEX coated PTNP The zeta potential of PTNP was found to be -17 ± 1.5 mV, similar to other DEX-coated nanoparticles (Figure 1D) (58, 59). The hydrodynamic diameters of nanoparticles were found to be larger than the diameters measured from TEM images due to coating and hydration layers (Figure S2B) (60). The hydrodynamic diameters of gold seeds, gold cores, core@shells and PTNP were found to be 18.2 ± 0.2, 79.2 ± 1.1 nm, 88.1 ± 2.2 and 87.7 ± 1.4 nm, respectively. In this study, DEX-coated PTNP were stable and no change in color was observed after 15 days’ incubation of nanoparticles with media, PBS, or saliva (Figure S3). Figure 1E shows the size distribution of PTNP measured with DLS for a week, which does not change over time, confirming the stability of these structures in deionized (DI) water.

### Optical behavior

Spherical gold nanoparticles (AuNPs) produce plasmonic properties and have been studied as potential contrast agents and anti-biofilm agent in different biomedical applications (40, 61), but they do not provide strong absorption spectra in NIR (Figure S4). Plasmonic properties in NIR are needed to make them an effective candidate in biomedical applications (56). Therefore, engineering AuNPs characteristics such as shape, size, and composition could help to establish a novel agent with the specific purpose of biofilm treatment methods. The plasmonic properties of shape-engineered gold in a gold cage nanoparticle were measured using UV-vis spectroscopy and are shown in Figure 1F. These nanoparticles have a plasmon peak at 808 nm which is tunable in NIR region based on its GRR and the thickness of shells (61). The application of these novel morphologies in photothermal therapy and photoacoustic imaging is an innovative approach to overcoming biofilms generated on tooth and wound surfaces.

### PTNP incorporation within biofilms

To measure the incorporation of PTNP within biofilm, different concentrations of PTNP (0, 0.1 and 0.25 mg ml^-1^) were coated with three different coatings, i.e. polyethylene glycol (PEG), dextran-10kDa (DEX) and polydopamine (PDA). **Figure S5** shows DEX enhanced incorporation of PTNP into S. mutans biofilms much more than PDA and PEG. We found that DEX coated PTNP were incorporated into S. mutans and S. aureus biofilms while using a low concentration of PTNP (0.25 mg ml^-1^). SEM images of biofilms incubated with PTNP indicated that PTNP bind to S. mutans biofilms (Figure 2A and B), which was confirmed by EDS measurements (Figure S6A-C). To investigate PTNP binding in biofilms of S. aureus we also carried out TEM of biofilms (Figure 2D and E), which showed clear uptake and penetration of PTNP within the biofilm ultrastructure. Figures 2C and F show that DEX-coated PTNP are significantly taken up into S. mutans and S. aureus biofilms as measured by ICP-OES. These data confirm DEX enhanced incorporation of PTNP into S. mutans and S. aureus biofilms, while no binding enhancement was observed on our control sample (saliva-coated apatite surface (sHA)). This polymer promotes selectivity towards biofilms while avoiding binding to host cells. Dextran is a component of biofilms and prior evidence suggests that dextran coated nanoparticles are incorporated into biofilms by the enzyme beta-glucosyltransferase (gtfB) (4, 62). These results demonstrate the successful incorporation of PTNP into S. aureus biofilms when applied topically, supporting the idea of using PTNP for photoacoustic imaging and photothermal therapy of biofilms.

**Figure 2.**
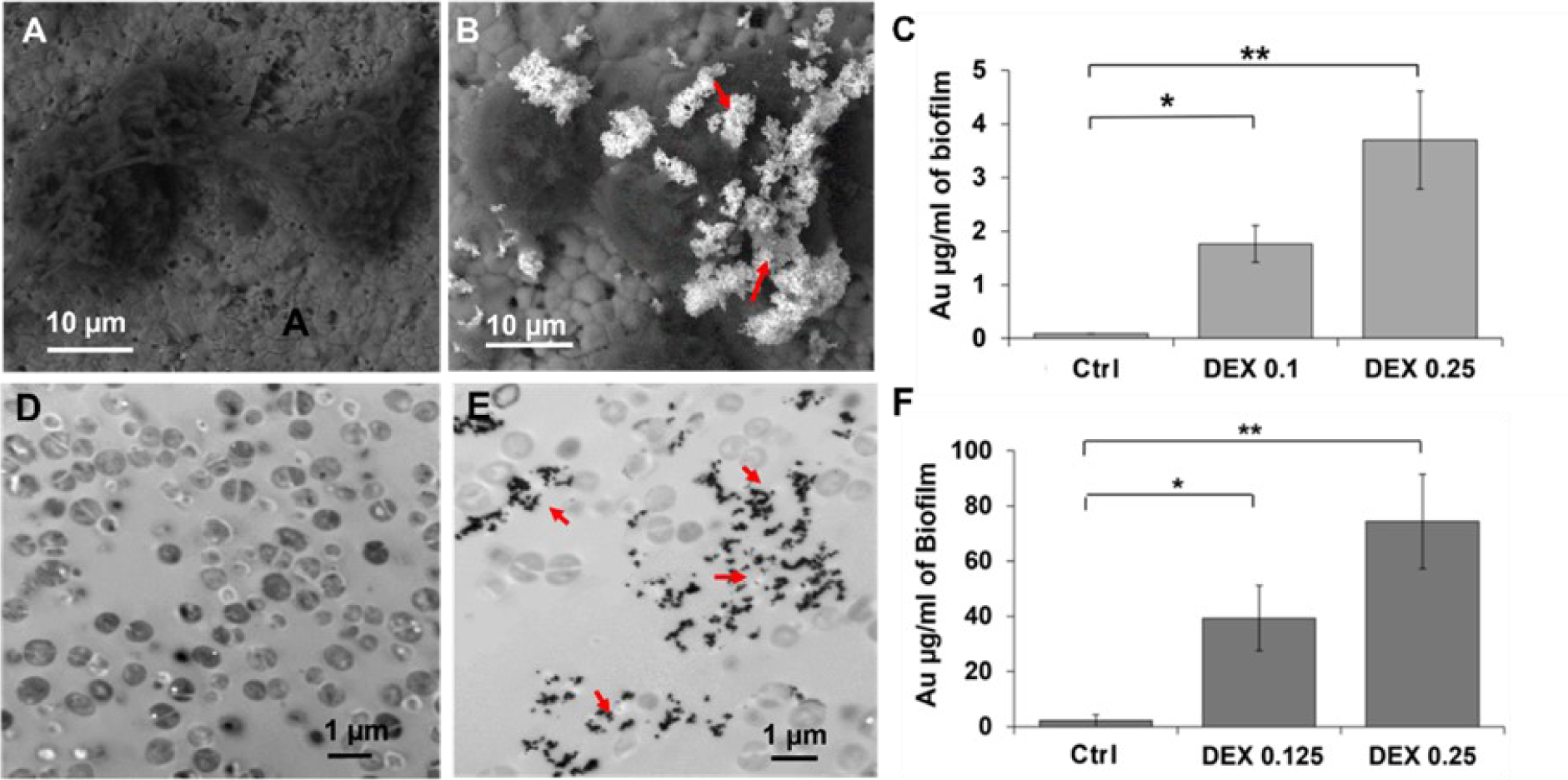
PTNP are taken up by bacterial biofilms. SEM in backscattered electron (BSE) mode showing the morphology of (**A**) untreated biofilm, (**B**) PTNP-treated S. mutans biofilm (Scale bar: 10 µm). (**C**) ICP-OES measurements on S. mutans biofilms. TEM of S. aureus biofilm uptake of PTNP, (**D**) untreated biofilm and (**E**) S. aureus biofilm treated with PTNP for 24 h (Scale bar: 1 µm). F) ICP-OES measurements on S. aureus biofilms. Red arrows show nanoparticles. (n=6), *P< 0.05, **P<0.005. The data were presented as mean ± SD.

### Photothermal activity of PTNP

Next, we examined the heat generation of PTNP while irradiated with NIR laser. PTNP solutions and DI water as a control solution were treated with NIR laser, and the temperature was recorded over time. PTNP structures were found to generate significant temperature increases, higher than the control solution. Figure 3A shows the maximum temperature observed for the PTNP solution was much higher than that for DI water (as much as 50 °C). The photothermal conversion efficiency of PTNP was calculated to be 77%, which is comparable with gold nanostars (78%) (63), gold nanoporous nanoshells (75.5%) (64), and higher than gold nanocages (53.6%) (65, 66), gold nanorings (42%) (67), gold nanoshells (41.4%) (68), gold nanorods (21.3%) (68, 69). The temperature change of the PTNP and its dependency on laser power and PTNP concentration was also studied. As is shown in Figure 3B, greater laser power density resulted in greater temperature increases. Similarly, increases in the concentration of PTNP led to greater temperature enhancement (Figure 3C).

**Figure 3.**
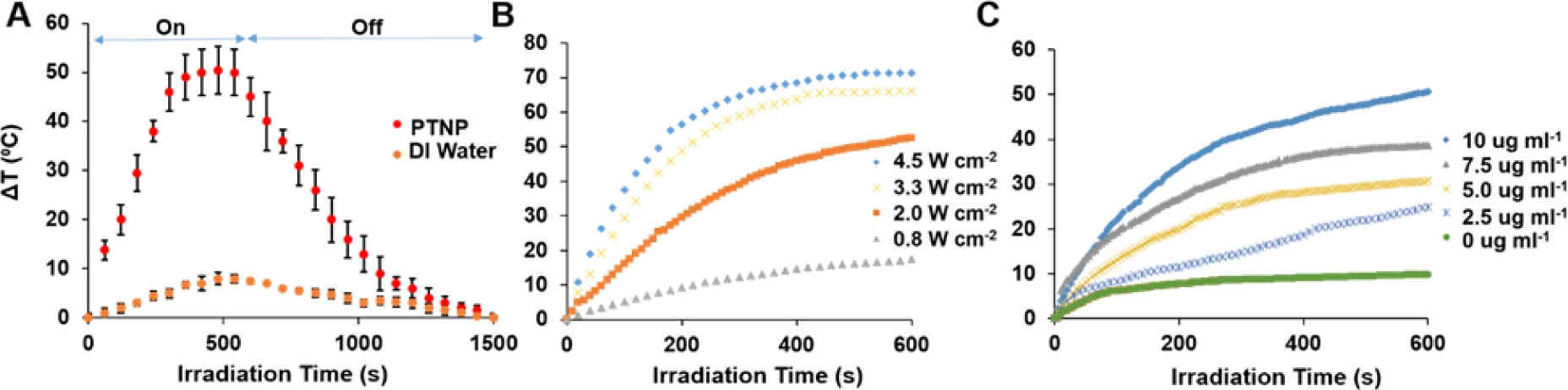
PTNP have excellent photothermal properties. (**A**) Heating and cooling curves of PTNP (10 µg ml^-1^) and DI water. (**B**) Temperature changes of solutions containing 10 µg ml^-1^ PTNP irradiated at laser power densities of 0.8, 2, 3.3 or 4.5 W cm^-2^. (**C**) Temperature changes of solutions containing 0, 2.5, 5, 7.5 or 10 µg ml^-1^ PTNP irradiated at a laser power density of 2 W cm^-2^ (n=5). The data were presented as mean ± SD.

We found that during five cycles of laser on-off irradiation, the PTNP solution reproducibly reached 99.9% of the temperature achieved the first time, indicating the excellent stability of these structures (Figure S7A). The TEM of irradiated PTNP samples do not show any difference in morphology, which confirms the photostability of these structures (Figure S7B and C). These results suggested that PTNP is suitable for photothermal applications. Here, we chose different laser powers depending on the experiment setting and in agreement with the work of others (67, 70, 71) For only in vitro and ex vivo oral biofilms that develop upon hard surfaces such as hydroxyapatite discs or teeth we used higher laser power (2 W cm^-2^) and shorter time frames which is again in agreement with recent studies (which report powers of 2 to 2.5 W cm^-2^) (71, 72). For in vitro and in vivo skin studies we used lower laser powers (0.25-0.7 W cm^-2^) to avoid any possible damage in surrounding tissues as has been reported in other studies (which use powers such as 0.75, 0.78, 1 W cm^-2^) (63, 73, 74).

To test the biofilm imaging ability of PTNP, we examined the photoacoustic (PA) signal of PTNP at different PTNP concentrations (0, 0.06, 0.12, 0.25 and 0.5 mg ml^-1^). Increasing PA signal was observed with increasing the concentration of PTNP (Figure S8 A, B). We also examined the PA properties of PTNP when incubated with S. aureus biofilm. The PA signal was assessed using different PTNP concentrations (0, 0.12, 0.25, and 0.5 mg ml^−1^) which were incubated with biofilm for 24 h and were imaged at a laser wavelength of 800 nm (Figure S9). The PA signal rose with increases in the concentration of PTNP. These findings suggest that PTNP can be used as an agent for both diagnosis and treatment of oral and skin infectious diseases.

### In vitro anti-biofilm efficacy of the PTNP

Here, we examined how PTNP combined with laser irradiation disrupts biofilms and kills bacteria. Oral biofilms (S. mutans UA159, a strain associated with endocarditis) were formed on sHA discs while S. aureus biofilms were grown in 96-wall plates as a skin infection model. Biofilms were analyzed using multi-photon confocal microscopy, colony counting (presented as colony forming units or CFU) and computational analysis (COMSTAT and Image J). Biofilms treated with or without PTNP (0.25 mg ml^-1^) were exposed to NIR laser irradiation at 808 nm in both wide area and precise spatial control irradiation mode (schematically shown in Figure 4A and B respectively), at 0.5 and 2 W cm^-2^ for different time points. Then, bacterial survival studies were conducted on PTNP surfaces using a colony formation assay both with and without NIR irradiation in a wide area mode. The biofilms were removed, homogenized, and the number of viable cells were determined.

**Figure 4.**
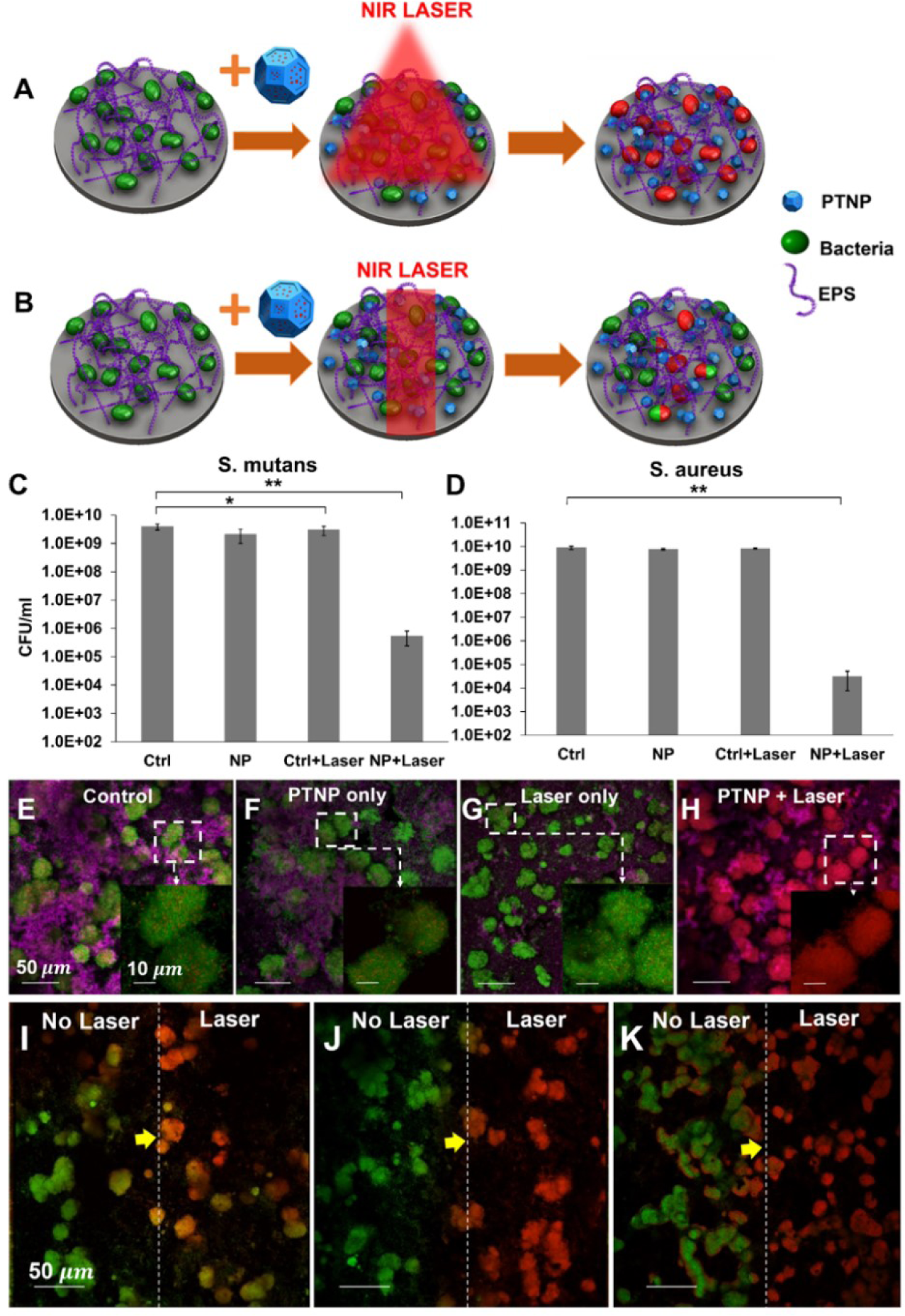
PTNP efficiently kill biofilm bacteria in vitro. (**A**) Schematic of photo ablation of biofilm with PTNP in wide area mode and (**B**) precise spatial control mode. The viability of (**C**) S. mutans and (**D**) S. aureus within biofilms treated with PTNP and irradiated in wide area mode (n=6). *P < 0.05, **P < 0.0001. (**E-H**) Representative confocal micrographs of treated and untreated S. mutans biofilms exposed to wide area mode (Scale bar: 50 µm). Magnified views of S. mutans biofilms in the boxed areas are shown in the bottom left of each panel (Scale bar: 10 µm). Confocal micrograph images of PTNP-treated S. mutans biofilms exposed to laser in precise control mode for (**I**) 15 s, (**J**) 30 s and (**K**) 60 s. Live bacteria cells (green), dead cells (red) and EPS (purple) are shown (Scale bar: 50 µm). Biofilm bacteria were irradiated at 808 nm (C-H and I-K were irradiated with laser power 0.5 W cm^-2^ and 2 W cm^-2^ respectively).

We observed an exceptionally strong biocidal effect against S. mutans and S. aureus within biofilms when exposed to NIR laser irradiation and PTNP, causing a >5-log and >6-log reduction (elimination of 99.99% of bacteria) of viable oral and skin pathogens, respectively (Figure 4C and D). This treatment is considerably more effective than the currently clinically used oral and skin antimicrobials such as chlorhexidine, alcoholic chlorhexidine, and 0.5% chlorhexidine-10% povidone iodine, which causes around 3-log reduction in same conditions (75–77). We observed that the dry weight of remaining S. mutans biofilms in the treated samples with NIR light was 41.1% less than the control samples and untreated samples (Figure S10). This was surprising, given the short laser irradiation time of this procedure (78, 79). These results suggest that the treatment can cause EPS degradation as well as killing the bacteria. These studies demonstrated that PTNP, when illuminated with NIR light, could effectively kill S. mutans and S. aureus, as well as degrading biofilm EPS.

### The effects of PTNP and laser treatment at a local and wide area level

To further understand how PTNP can be used as a biofilm treatment, we performed treatment experiments using confocal microscopy. The same treatment procedure was repeated (irradiating the samples with NIR laser in wide area mode, schematic is shown in Figure 4A) in the last section. Representative images of control, PTNP-treated biofilms, exposed control biofilm and PTNP-treated biofilms to the NIR laser are shown in Figure 4E-H. We observed almost complete bacteria killing in biofilms incubated with PTNP and after 10 min of wide area mode laser treatment, while few dead cells were observed in other samples. These results confirmed our hypothesis that photothermal ablation PTNP can kill S. mutans bacteria.

To underscore the potential targeted approach of our therapy, we used a modified version of the treatment method described above. Here, the laser beam was focused to an area in the middle of the biofilm covered disc of 2 mm x 5 mm and turned on for 15 s, 30 s or 60 s (Figure 4J-K). The untreated area is on the left and the treated area on the right in Figure 4I-K. The 15 s NIR laser irradiation is not enough to completely kill the bacteria in the irradiated area; however, 30 s irradiation can completely kill the bacteria, and there is a clear line between live and dead cells (the yellow arrows show the line). The 60 s exposure resulted in bacterial killing even in the unirradiated part of the biofilm, which indicates that for in vitro localized treatment, 60 s irradiation is too long. Here, we demonstrate how our novel anti-biofilm agent could treat the bacteria locally in less than a minute.

### Biocompatibility and cell toxicity of PTNP

In vitro biocompatibility and determining the photothermal activity of PTNP were carried out on different cell types. The cells were treated with DEX-coated PTNP for 4 hours at concentrations of 0.1, 0.25, and 0.5 mg ml^-1^ PTNP. The viabilities of the C2BBe1, BJ5IA, gingival and HaCaT cell lines were determined using the MTS assay. It was found that these structures did not significantly affect the viability of these cell types and are biocompatible under the conditions tested (Figure 5A).

**Figure 5.**
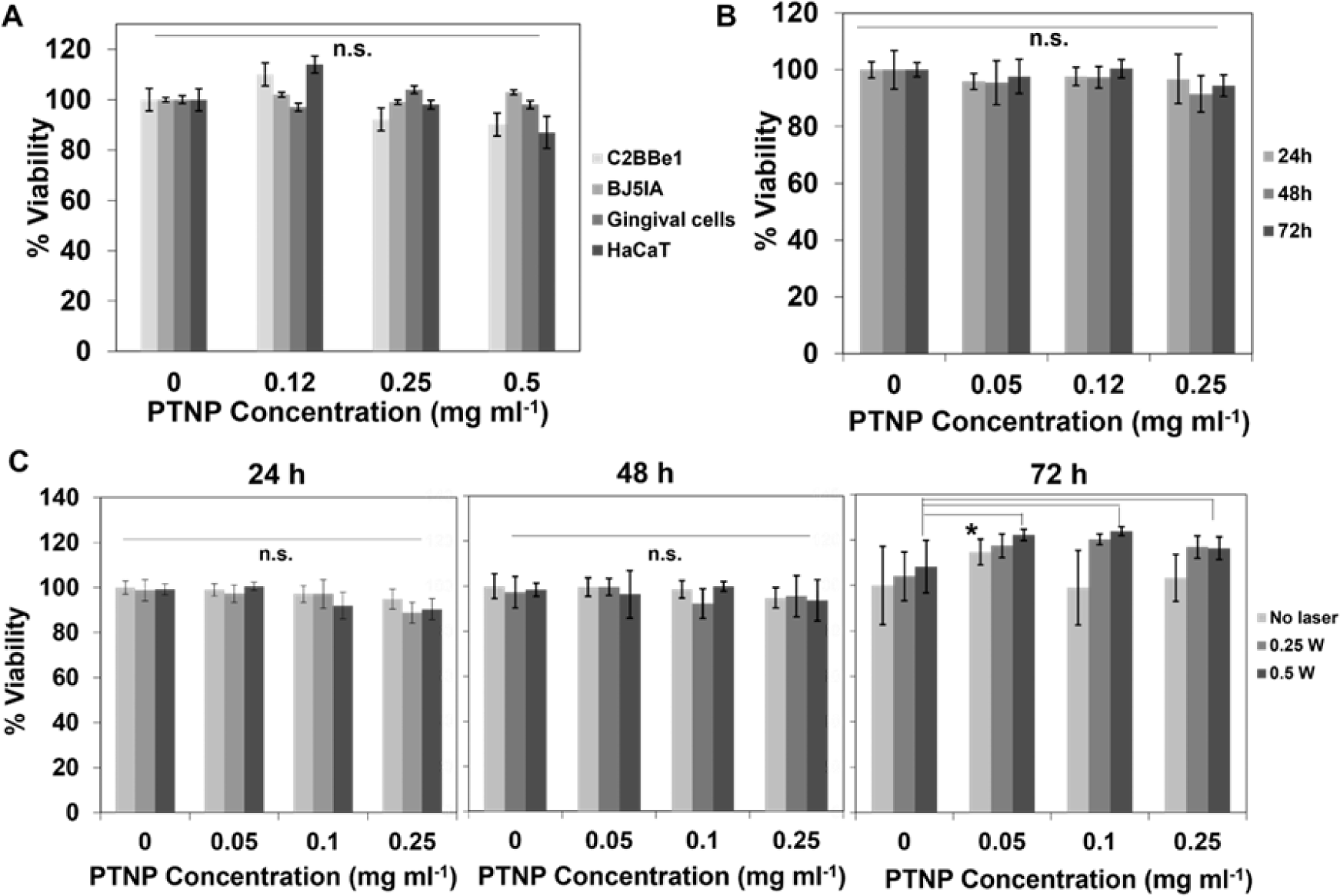
PTNP are biocompatible and non-toxic. (**A**) The effect of PTNP on the viability of several cell lines. (**B**) The cell viability of HaCaT cells in 24, 48, and 72 h of incubation with PTNP. (**C**) Cell cytotoxicity of HaCaT cells incubated with PTNP in different concentrations (0.1, 0.05, 0.1 and 0.25 mg ml^-1^) and time points. Cells were irradiated at 808 nm (0.25, and 0.5 W cm^-2^). (n=3), * = P < 0.05. “n.s.” = not significant. Error bars are standard deviations. The data were presented as mean ± SD.

To evaluate the precision and selectivity of this treatment method for bacterial cells, we evaluated whether this approach could target biofilms without adversely impacting surrounding tissue. Since the amount of heat generated by PTNP can be controlled by either changing the laser power or concentration of PTNP in the medium, it can provide fine control and homogenous distribution of heat compared to conventional heating probes (80). For these experiments, HaCaT cells were selected and incubated with different concentrations of PTNP (0.05, 0.12, and 0.25 mg ml^-1^) without light, and their viability were quantified after 24, 48 or 72 hours (Figure 5B). These figures confirm the biocompatibility and nontoxicity of PTNP within different cell lines. In this study, PTNP has been used as an antibiofilm for topical applications where scant uptake is to be expected, and therefore persistence and lack of ability to biodegrade is of lesser concern. In parallel experiments, treatment with PTNP was combined with laser irradiation at different powers (0.25 and 0.5 W cm^−2^, 7 min) and HaCaT cell viability was tested at 24, 48, or 72 h after laser irradiation (Figure 5C). Live-Dead staining microscopy image of HaCaT cells at 0.5 W laser irradiation at different time points is shown in Figure S11. We found that PTNP is well-tolerated by eukaryotic cells and ineffective at killing them via laser-irradiation due to lower uptake (Figure S12) and greater robustness compared to bacteria. Moreover, the results indicate regrowth of the cells after 72 h of laser irradiation, which shows that the effects of laser on mammalian cells were negligible, and no cell damage was observed on NIR irradiated PTNP-treated cells at different powers of 0.25 and 0.5 W cm^−2^ for 7 min. This study also suggests the feasibility of carrying out photothermal anti-biofilm treatment using PTNP in both oral and wound in vivo models, while keeping the surrounding tissue healthy.

## PTNP effects in animal models

### PTNP detects and kills pathogenic oral bacteria in an animal model

The anti-bacterial efficacy of PTNP, with a clinically relevant topical treatment regimen for oral infections, was examined using Sprague-Dawley rats infected orally using an actively growing culture of S. mutans (81). Ex vivo PTT efficacy and PA experiment were performed on rat’s teeth as is shown schematically in Figure 6A. PTNP nanoparticles (0.25 mg ml^-1^) were applied topically on the teeth and after 10 min incubation within biofilms, they were irradiated with the NIR light for 1 min. PTNP absorbs the light and generates local high temperatures that kill S. mutans bacteria. Figure 6B shows the viability of S. mutans within PTNP treated biofilms after the laser treatments. Here we found that none of the other groups (control, only PTNP, only laser and Chlorhexidine) are able to kill the bacteria successfully, and our developed treatment method was more than 99.99% effective in killing the S. mutans bacteria. To evaluate the effectiveness of laser treatment in situ, the viability of bacterial cells within treated biofilms was analyzed by confocal laser scanning microscopy (Figure 6C). As shown in Figure 6C, the biofilm bacteria are alive after treatment in control, PTNP only and laser only groups. But the PTNP with laser treatment resulted in highly effective method which rapidly killed most of the bacteria. We also noticed that while the biofilm treated by chlorhexidine harbored predominantly dead bacteria in imaging, the bacteria recovered from the jaw was not significantly reduced. This may reflect differences in the location of the imaging analysis and recovery of plaque from both smooth and sulcal surfaces for the CFU counting. For confocal imaging, given the anatomical constraints of the jaw, we could only visualize the live/dead bacteria on smooth surface of the teeth, which could not depict the large number of live bacteria inside the sulcus that is more difficult to kill. Chlorhexidine, a highly cationic molecule, is known for its poor penetration into deeper biofilms. As a result, the bacteria in the deep layers (e.g., sulcal areas) might remain alive while most of the bacteria on smooth surfaces would be killed (3). For CFU counting, the jaws were subject to an optimized sonication procedure that recovers all the bacteria from both smooth and sulcal surfaces, which are harvested and counted. Despite technical differences, both confocal imaging and CFU counting clearly show that PTNP with laser were highly effective against the plaque bacteria compared to other conditions.

**Figure 6.**
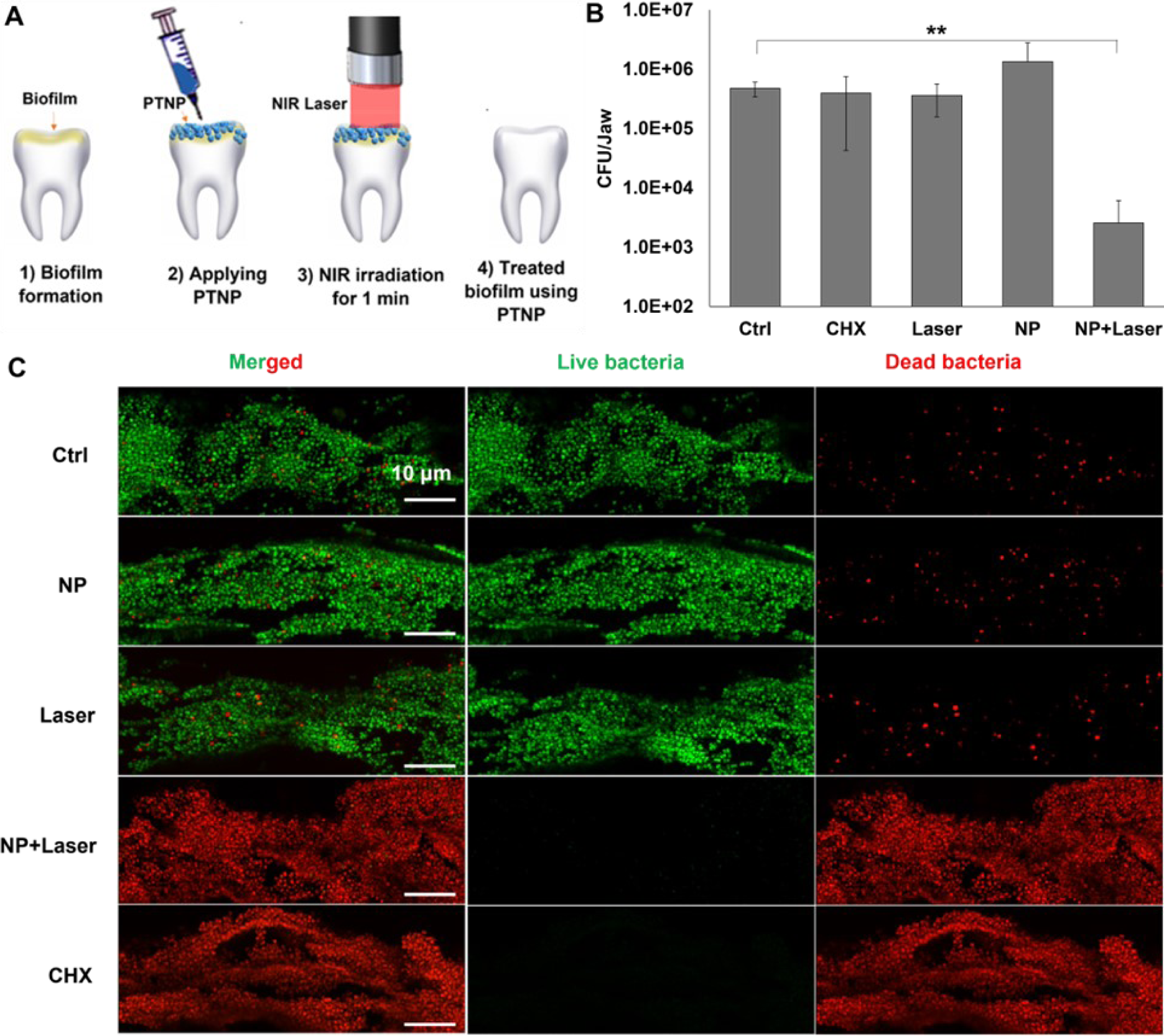
PTNP are multi-functional nanoagents that detect and treat oral infections. Photothermal efficacy of PTNP in the oral model; (**A**) A schematic of the PTT experiment. (**B**) Bacteria killing in an animal model of oral disease, as evidenced by the viability of S. mutans (n=6). (**C**) Representative confocal microscopy of control, PTNP only, Laser only, PTNP and laser-treated, and chlorhexidine only within S. mutans biofilms. Live and dead bacteria are shown in green and red respectively (Scale bar: 10 µm). ** indicate statistically significant differences at P<0.005.

We next evaluated biofilm imaging and diagnostic potential of PTNP in our oral model. For the oral PA experiment, we imaged jaws incubated with either vehicle (a buffer or solvent as a control) or PTNP (Figure 7A and B). The PA images reveal strong contrast for the teeth which had been treated with PTNP, while no signal was detected for the control group (Figure 7C and D). PA image analysis showed that infected rat teeth that were incubated with PTNP have a 72- fold signal increase compared to the control (Figure 7E). Photoacoustic imaging has recently been used in oral applications in patients (49, 82) and these data confirm the infection on the teeth using PTNP and suggests that PA could also be used to diagnose oral infections.

**Figure 7.**
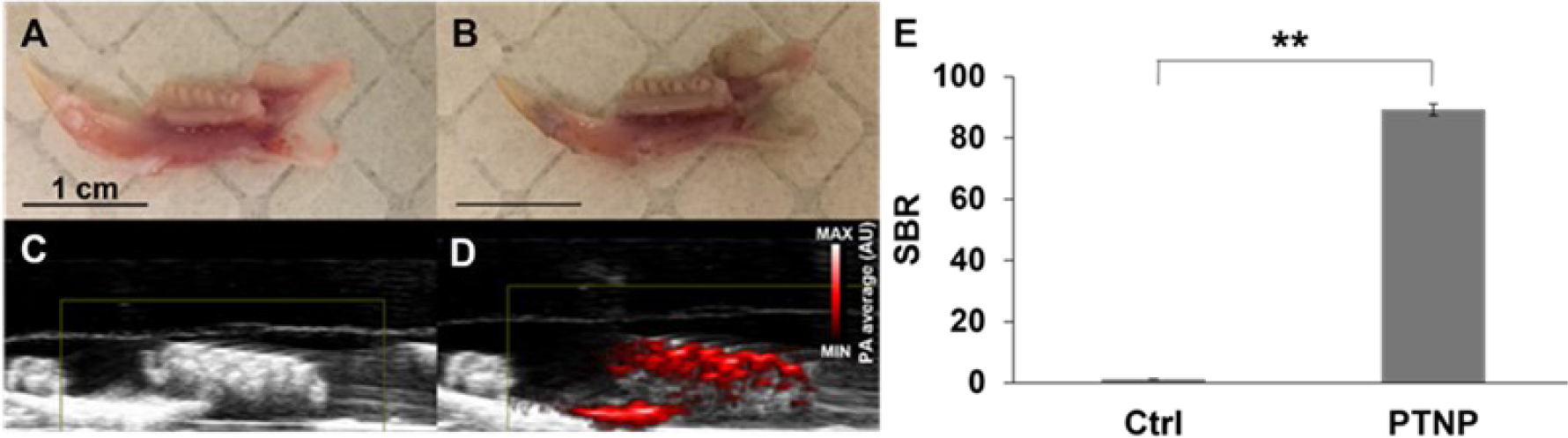
PTNP exhibit strong photoacoustic imaging signal in our oral model. Representative photographs of infected teeth (**A**) control, (**B**) incubated with PTNP (Scale bar: 1 cm). Grayscale ultrasound images overlaid with PA images of (**C**) control (**D**) incubated with PTNP. (**E**) SBR of the infected teeth with and without PTNP (n=6). ** indicate statistically significant differences at P<0.005.

### Treatment of Staphylococcus aureus infected wounds using PTNP anti-biofilm agents

The feasibility of this treatment strategy to disinfect wounds was also investigated in vivo. A delayed inoculation S. aureus wound model was used to test the applicability of this technique (83, 84). As presented in Figure 8A, infection progress was monitored using bioluminescence imaging (BLI). The PTNP with laser group was the only group that showed no infections in the BLI images after 5 min of treatment (Figure 8B). To confirm the reliability of this novel technique, animals were euthanized, and wound tissues were extracted for bacterial plate counting (85–87).

**Figure 8.**
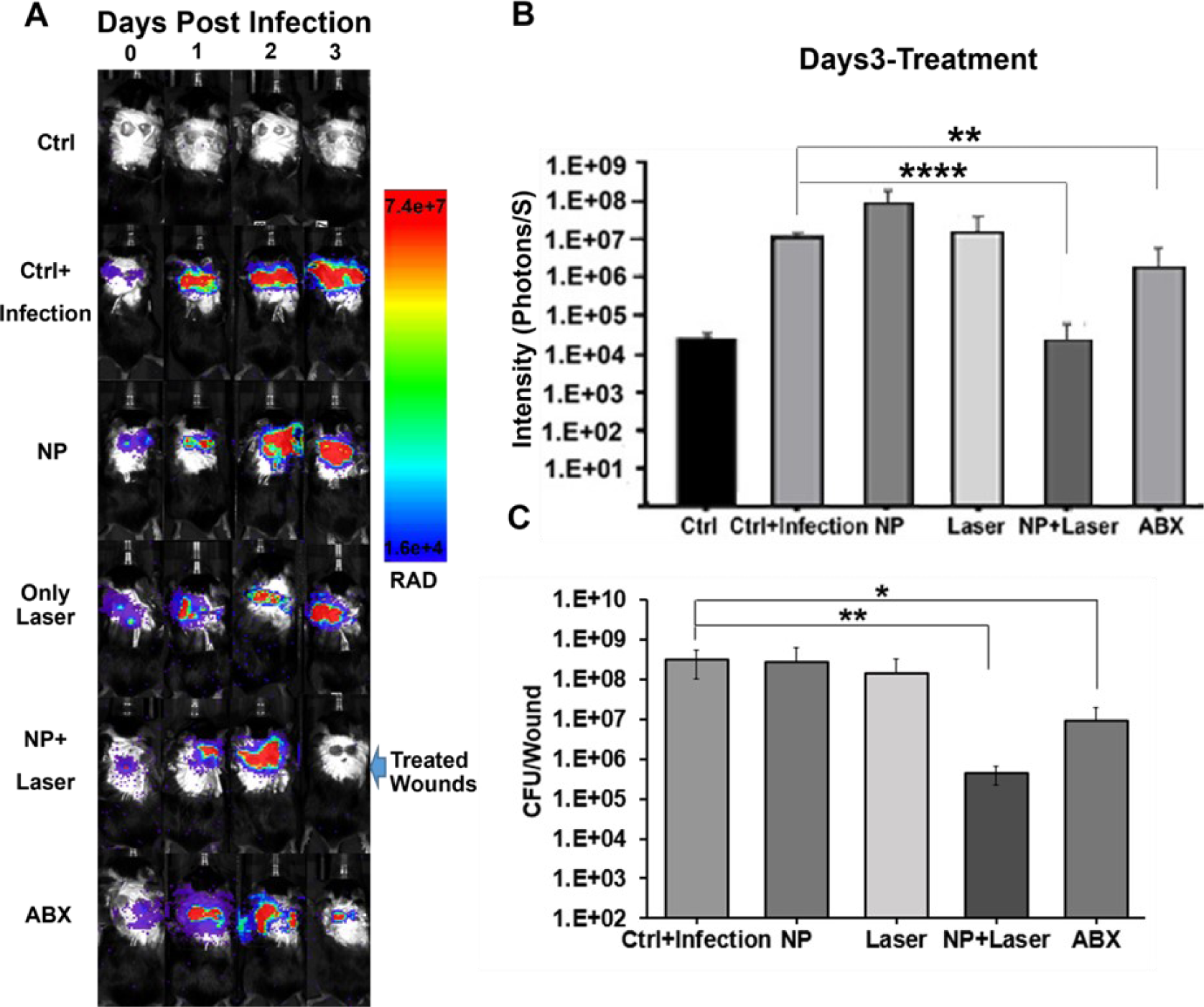
Bioluminescence in vivo imaging of an excisional wounds’ infection model confirms the efficacy of PTNP. (**A**) BLI of mice infected with S. aureus. (**B**) Analysis of bioluminescence signal intensity after treatment. (**C**) The viability of S. aureus extracted from wounds treated as noted. In the graphs Ctrl and NP denote control and PTNP. (n=6), *, **, and **** indicate statistically significant differences at P<0.05, P<0.005, and P<0.0005.

Figure 8C shows the viability of S. aureus in our wound model for different treatment groups. Infected wounds which were treated with PTNP and laser irradiation had a remarkable decline in bacteria viability, in agreement with our in vitro studies. These results suggest that the treatment of the infected wounds with S. aureus bacteria using PTNP and laser killed most of the bacteria within a short time, however the laser only, PTNP only and topical gentamicin, an antibiotic positive control (ABX), did not have the same killing effect.

One of the greatest advantages of this method is to be able to control the heat generated by PTNP and monitor the infected wounds using an infrared thermographic camera. Here, the camera detected the surface temperature of tissues, treated and non-treated wounds. We observed an increase in wound temperature for the groups treated with PTNP during laser irradiation (Figure S13A and B). However, ABX, control, laser, and PTNP only treated mice showed the least heating effect, increasing approximately 5 °C above mice’s body temperature; this observation is in line with our in vitro data showing an increase in temperature of DI water when irradiated with NIR laser (Figure 3A). On the other hand, wound temperature increased by 20.3 ± 1.4 °C when treated with PTNP and laser irradiation for 5 min (Figure S13C). The topical treatment of infected wounds using this novel method provides the ability to control heat delivery at a local scale. As demonstrated in Figure 4A and B, depending on the irradiation strategy used (i.e., whether a diffuse or focused laser beam), homogenous or localized biofilm treatment can be achieved that effectively kills bacteria without damage to the surrounding tissues in a short treatment duration. Here, we adjusted the NIR light to irradiate only the infected parts. This approach represents a potentially valuable treatment method in which an optimized nanoparticle can be employed for biofilm- associated infectious diseases treatment such as catheters, heart valves, and prosthetic joints.

We also examined our multi-functional nanoagent to image wound infections in vivo. To visualize the infected wounds, 20 μL of 2.5 mg ml^−1^ of PTNP solution was dropped onto the wounds area. Then, the PA imaging was performed after 10 min, while the mice were still under anesthesia. Similar to oral biofilms, PTNP-treated mice showed high signal intensity localized to the infectious wounds, while no signal was detected in the controls (Figure S14).

## Discussion

Here, we have demonstrated that PTNP are an effective theranostic anti-biofilm agent, enabling rapid photoablation and outstanding biofilm disruption. We show the utility of approach in vitro, against two different bacterial species, and in a pair of in vivo preclinical models. This approach represents advancements over existing theranostic agents in demonstrating strong extinction spectra in the NIR region, excellent photothermal conversion efficiency, capability in imaging of biofilm bacteria and eliminating them rapidly. A variety of gold nanostructures have been used for PTT previously, such nanorods and nanoshells. However, the synthesis of those structures typically involves the use of organic solvents or toxic chemicals, which require cumbersome procedures to eliminate or displace. The synthesis of PTNP, as described herein, avoids the use of such organic solvents and toxic reagents, with the primary reagents used being safe materials, i.e. sodium citrate and ascorbic acid. Therefore, the prospects for scale-up synthesis and clinical translation of PTNP are improved versus nanorods and nanoshells (65, 66, 68).

PTNP demonstrates several remarkable features that are useful for the treatment of infectious diseases. First, it has a unique structure and is non-toxic and biocompatible, which provides considerable interest for biomedical applications. Second, it can be used as a dual-use agent for both imaging and therapeutic applications to diagnose, degrade, and disinfect oral and skin biofilms. Third, it can be used in both wide-area irradiation and precise spatial control, which could provide homogenous or localized biofilm treatment to effectively target, activate and kill bacteria. Fourth, this treatment method using PTNP is antibiotic-free, and therefore there is no possibility of antibiotic resistance developing. Fifth, the biofilm killing using this approach is expeditious and low cost. We find that the amount of the bacteria decreases dramatically by using 3.75 mg of PTNP (when scaling to a human dose of 20 ml), which will only cost about $0.20. The other equipment that is needed to perform the biofilm treatment is a NIR laser that can be used for many years and costs around $200, which is an affordable expense for a clinical office. Alternatively, these agents could be used by patients at home, in combination with low cost, whole surface irradiators, which are already on the market. Our vision is that these PTNP could be incorporated into a variety of daily oral and skin health maintenance products, such as mouth rinses, gel strips, and photothermal bandages.

This study has some limitations. We have assessed biofilm killing but not therapeutic outcomes (e.g., preventing cavities or enhancing wound healing, so future studies will investigate whether PTNP can control cavities or accelerate wound healing). We tested our anti-biofilm agent with a single wavelength NIR light. This can be fine-tuned further or tested on other light sources. Exploring drug loading or targeting within PTNP structures to either enhance anti-biofilm efficacy or other functionalities (e.g., growth factors, anti-demineralizing agents) could also lead to improved therapeutic outcomes. Moreover, the use of this treatment method and anti-biofilm agent for other biofilm infections including those in catheters, heart valves, lungs, and prosthetic joints could be investigated.

In summary, dual purpose biocompatible PTNP structures, which have photothermal characteristics that detect, target, and kill bacteria within biofilms without the need for broad-spectrum antibiotics, rapid effect etc., is highly appealing for oral and skin infectious diseases. These properties allow the development of a range of different oral care products and wound disinfections. Here, we showed that this treatment method can accurately and selectively deliver the desired amount of heat to the infected area without damaging the healthy tissue and can be carried out in a very short time to target infected bacteria. The results of this research will advance the field of biofilm treatments by introducing novel structures of anti-biofilm agents that are activated under illumination by NIR light, resulting in outstanding biofilm disruption in vitro and in vivo in less than a minute.

## Materials and Methods

### Nanomaterial synthesis and characterization

PTNP were synthesized via the Turkevich method, a seeded growth method, and GRR using chloroauric acid (HAuCl_4_). In brief, 16 ± 2 nm seeds were prepared using the Turkevich method, and 78 nm cores were synthesized using the seeded growth method. 3 ml of 1 mM aqueous AgNO_3_ (Sigma-Aldrich) was reduced on the 78 ± 3 nm cores using 1 ml of 1 mM L-ascorbic acid (AA) (Fisher Scientific), making core@shell structures. Then, 10 ml of 500 μM aqueous HAuCl_4_ (Sigma-Aldrich), was added at a rate of one drop per second to create Au@Au core-cage structures, here termed PTNP (56). Ligand exchange was performed using 5 ml of 0.1 M dextran solution (Dextran T-10, Pharmacosmos). Next, the mixture of PTNP and dextran solution was allowed to stir overnight. The final, coated nanoparticles were then collected by centrifugation at 1200 rcf for 10 min.

### UV-visible spectroscopy

UV-visible spectra were recorded from wavelengths of 300 to 1000 nm and obtained from 1 ml of diluted PTNP using a UV-visible spectrophotometer (Thermo-Fisher Scientific).

### Electron microscopy

Morphological and structural characterization of PTNP was done determined using a transition electron microscope (TEM, JEOL 1010) operating at 80 kV and scanning electron microscope (SEM, Quanta 600 FEG, FEI). We examined PTNP binding using TEM to visualize and analyze PTNP distribution within skin biofilm ultrastructure (88). First, S. aureus biofilms treated either with PTNP or vehicle (Figure S3) were washed with DI water to remove unbound materials. Biofilms were fixed with 2.5% glutaraldehyde and 2.0% paraformaldehyde in 0.1 M sodium cacodylate buffer, pH 7.4, overnight at 4 °C. Prepared samples were fixed in 2.0% osmium tetroxide with 1.5% K_3_Fe (CN)_6_ for 1 h at room temperature and rinsed in DI water. Then, samples were dried and were embedded in EMbed-812 (Electron Microscopy Sciences). 1% uranyl acetate and SATO lead were used to stain the thin sections of the final samples. TEM images acquired using a JEOL 1010 electron microscope fitted with a Hamamatsu digital camera and AMT Advantage NanoSprint500 software. In a separate experiment, PTNP binding to intact biofilms was also assessed. Briefly, biofilms vehicle treated and PTNP-treated biofilms were dip-washed three times with 0.1 M NaOAc buffer (pH 4.5) and transferred to the 0.1 M NaOAc reaction buffer (pH 4.5) containing TMB and H_2_O_2_. After 30 min, the intact vehicles treated biofilms and biofilms treated with PTNP were removed and were fixed with 2.5% (v/v) glutaraldehyde (GA) in PBS at 4 °C overnight. EDS was also performed to detect gold and silver from PTNP, with corresponding elemental mapping.

### Hydrodynamic diameter and zeta potential measurements

Hydrodynamic diameter and zeta potential of PTNP were examined using dynamic and electrophoretic light scattering (Nano ZS-90 Zeta sizer). All DLS data were acquired by diluting 100 µl of 0.1 mg ml^-1^ PTNP structures to 1 ml with DI water.

### Inductively coupled plasma optical emission spectroscopy

PTNP concentrations were measured using inductively coupled plasma optical emission spectroscopy (ICP-OES, Spectro Analytical Instruments GmbH) to measure the gold and silver concentrations in PTNP samples and also within tissue samples according to a protocol published elsewhere (56).

The total amount of gold within intact oral and skin biofilms was also measured using ICP-OES. Biofilms treated with PTNP were transferred to glass tubes and digested with 1 ml of aqua regia for 1 h at room temperature and then centrifuged at 1000 rpm for 5 min to remove any excess supernatant. Then, the volume was adjusted to 10 ml with DI water prior to analysis with ICP-OES. For these experiments, six samples were prepared for each PTNP, and the data were presented as mean ± SD.

### In vitro heating

The photothermal properties of PTNP was measured using NIR laser irradiation to induce heating of these structures. 1 ml of PTNP (10 µg ml^-1^) was added to a 2 ml quartz cuvette, and a fiber optic thermometer (Nomad, Qualitrol-Neoptix) was inserted 5 mm below the liquid surface. Solutions were then irradiated with an 808 nm laser (OEM Laser Systems) at 2 W cm^-2^ power for 10 min. Temperature was recorded every 10 s. DI water was used with the same procedure as a control. For the laser power and PTNP concentration dependency experiment, the laser was turned off after reaching a constant temperature (i.e. after 10 min). The photostability (Figure S5) was measured with 10 µg ml^-1^ PTNP and at NIR laser irradiation with 2 W cm^-2^ while the laser was turned off after reaching a plateau in the temperature, the sample was allowed to cool down to the room temperature, and the laser was again turned on to reach the maximum temperature for the next round. The final temperature is given as a delta to exclude minor variations in baseline temperature and to highlight the differences in temperature gain caused by the conditions used. This experiment was repeated 5 times. All experiments were performed at room temperature with ambient light.

### Laser Set-up for in vitro and in vivo experiments

Irradiation using collimated laser beam (Precise spatial control mode): The experiments in which light was collimated and the flux was precisely controlled have been carried out using NIR laser (808 nm, OEM Laser Systems, flux 0.5 or 2 W/cm^2^). The beam had a rectangular cross-section (2mm*8mm), and it could cover the entire teeth of a rat. The object was irradiated during different time periods (15 s, 30 s, 1 min, 5 min, and 10 min). This approach can be clinically translated and used for treatment of teeth cavities and biofilm-forming infections in humans.

Wide-area irradiation: The same laser coupled to a fiber (808 nm, OEM Laser Systems) could be used for wide-area irradiation. The beam was focused on the entrance aperture of a multimode glass fiber (Dolan Jenner, 1 m, 4 mm diameter) using a lens (f=5.5 mcm) mounted on a 1D translation stage (Thorlabs) such that the size of the spot on the fiber aperture was ∼2-3 mm in diameter. Such slight defocusing allowed us to avoid overly high fluxes at the aperture and prevent overheating. The light emerged from the other end of the fiber uncollimated, and therefore by adjusting the distance between the fiber tip and the object, round areas of different diameter could be irradiated. The photon flux across the area was approximately uniform, as judged by visual examination, and it was calculated from the laser power, measured by an optical power meter (Thorlabs) immediately after the fiber and the area diameter.

### In vitro oral biofilm model

Streptococcus mutans (S. mutans) UA159 (ATCC 700610) was used as an oral biofilm bacterium and was grown in ultra filtered (10 kDa molecular-mass cutoff) tryptone-yeast extract broth (UFTYE; 2.5% tryptone and 1.5% yeast extract) containing 1% (wt/vol) glucose at 37 °C and 5% CO_2_ to mid exponential phase. S. mutans biofilms were formed on saliva-coated hydroxyapatite (sHA) discs with surface area of 2.7 ± 0.2 cm^2^ (Clarkson Chromatography Products Inc., South Williamsport, PA), as described previously (85). Each HA disc was coated with filter-sterilized saliva for 1 h at 37 °C (the saliva was prepared as described elsewhere).(89) The discs were vertically suspended in 24-well plates using a customized wire disc holder that mimics the smooth surfaces of the tooth. These sHA discs were each inoculated with ∼ 2 × 10^5^ CFU of S. mutans per milliliter in UFTYE culture medium (pH 7.0) containing 1% (w/v) sucrose at 37 °C with 5% CO_2_. The culture medium was changed at 19 and 29 h (twice daily) until 43 h. Then, the biofilms were collected and analyzed for PTNP binding, biomass reduction and bacterial killing as described below.

### Bacterial killing and biomass reduction by PTNP with laser irradiation

To assess the anti-biofilm effect of PTNP within biofilms, the bacteria were grown overnight and then 1:100 dilute from overnight culture were added into fresh medium for biofilm assay. For the skin biofilm formation, S. aureus isolates (USA300, Newman 502A) were grown at 37°C in liquid Luria broth (Fisher Scientific) at 300 rpm. Then 100 μL of the dilution was added per well in a 96 well dish and the plates were incubated at 37°C with 5% CO_2_ for 24 h. Then the plates were washed 3 times with DI water before incubating with different concentrations of PTNP (0, 0.1, 0.25, and 0.5 mg ml^-1^) for another 24 h. For oral biofilm formation, the S. mutans biofilms were grown on sHA discs as described above. The sHA discs and biofilms were topically treated twice-daily by placing them in 2.8 ml of PTNP (at either 0.12 or 0.25 mg ml^-1^) in 0.1 M NaAc (pH 4.5) for 10 min at room temperature at specific time points (Figure S2). After incubating the discs with PTNP, all the discs (controls and PTNP samples) were inserted in DI water 3 times and washed out completely. At the end of the experiment, the PTNP-treated biofilms were exposed to NIR laser (808 nm, OEM Laser Systems, 808 nm at a power of 0.5 and 2 W cm^−2^) for different time periods (15 s, 30 s, 1 min, 5 min, and 10 min). After NIR laser exposure, some samples were used for visualization and confocal microscopy as described below, and some were used for CFU counting and dry weight assay. For the latter purposes, the biofilms were washed with sterile saline solution (0.89% NaCl) three times and removed by a spatula from sHA discs. Then, the samples were homogenized using bath sonication (90, 91). Samples of these biofilm suspensions were diluted in different concentrations and plated onto blood agar plates. Then, these plates were placed in an incubator at 37 °C with 5% CO_2_ for 48 h. The total numbers of viable cells in each biofilm control and treated biofilm were determined by counting colony forming units (CFU). The remaining suspension was centrifuged at 5500 g for 10 min, and then washed with DI water twice and dried in oven at 90 °C for 2 h. Lastly, the final residue was weighed to assess biomass reduction (13, 90).

### Dynamics of bacterial killing of PTNP within intact biofilm and EPS structures

The distribution of PTNP, the bacterial killing effect of PTNP and EPS degradation within biofilm were visualized using a confocal fluorescence microscope (LSM 800, Zeiss) with a 20× (numerical aperture, 1.0) water immersion objective. Live and dead cells were labeled using SYTO 9 (485/498 nm; Molecular Probes) and propidium iodide (PI, 535/617 nm; Molecular Probes), and Alexa Fluor 647-dextran conjugate (647/668 nm; Molecular Probes) was used for labeling EPS.

### Cell viability

BJ5ta (human fibroblast) and C2BBe1 (human colorectal adenocarcinoma, epithelial) cells were purchased from ATCC (Manassas). Primary human gingival epithelial cells and HaCaT were donated by Drs. Manju Benakanakere and John Seykora’s lab in Dental School and the Dermatology Department of University of Pennsylvania, respectively. The effect on cell viability of PTNP was assessed using the MTS [(3-(4,5-dimethylthiazol-2-yl)-5-(3- carboxymethoxyphenyl)-2-(4-sulfophenyl)-2H-tetrazolium)] assay (CellTiter 96 cell proliferation assay kit; Promega, WI, USA) (92). Dulbecco’s modified eagle’s medium (DMEM) and medium 199, 10% fetal bovine serum (FBS) (Gibco) and 0.01 mg ml^-1^ of hygromycin B (Sigma-Aldrich) were used as a culture medium for BJ5ta cells. C2BBe1 cells were also cultured in a medium of DMEM, 10% FBS, 45 IU ml^-1^ penicillin and 45 IU ml^-1^ streptomycin (Gibco), with 10 µg ml^-1^ human transferrin (Sigma-Aldrich). Primary human gingival epithelial cells were cultured in keratinocyte serum-free medium (Invitrogen) and HaCaT cells were cultured in DMEM with Glutamax,4.5 g/l D-glucose, 110 mg l^-1^ Na Pyruvate (Gibco/Invitrogen #10569), 5% FBS and mixed with antibiotics streptomycin (100 µg ml^-1^) and penicillin (100 U ml^-1^). The cells were grown at 37 °C in a 5% CO_2_ humidified incubator. To determine the cytotoxicity of PTNP, 10^4^ cells per well were seeded in 96- well plates, and then the plates were incubated at 37 °C in a 5% CO_2_ atmosphere for 4 h (the cell viability of HaCaT cells were also measured after 24, 48, and 72 h incubation with PTNP). Then, the media was carefully removed, and the cells were washed gently with sterile phosphate buffered saline (PBS) twice and replaced with fresh media containing PTNP at different concentrations. The plates were then incubated at 37 °C in 5% CO_2_ atmosphere for different times (24, 48, and 72 h). After this incubation, the media was removed, the cells were washed with PBS, then 20 μl MTS reagent and 100 μl cell culture medium was added to each well. The plates were incubated at 37 °C in a 5% CO_2_ atmosphere for 1 h, then the absorbance was measured at 490 nm using a plate reader. The percentage of relative cytotoxicity of each concentration of PTNP was calculated compared to control and the data presented as mean ± standard deviation (n = 3). The effect of PTNP on cell viability was also evaluated while the cells were irradiated with the NIR laser. In these experiments, HaCaT cells were plated in 24-well plates (10000 cells per well) and incubated at 37 °C in a 5% CO_2_ atmosphere for 24 h. Then, the cells were washed with PBS and exchanged with the fresh media containing PTNP with different concentrations (n = 3 wells per condition). The following day, the cells were washed with PBS twice and replaced with fresh media. Immediately after media exchange, wells were irradiated with a NIR laser (808 nm wavelength, 0.25 and 0.5 W cm^-2^) for 7 min. Plates were then incubated at 37 °C in a 5% CO_2_ atmosphere to be analyzed for laser cytotoxicity of cells at 2, 24, 48, 72 h after laser irradiation using the Live-Dead assay. For this purpose, after reaching each time point, the cells were washed twice with PBS, and the media was exchanged with the Live-Dead reagents consisting of 2 ml PBS, 2 µl ethidium-1 homodimer for dead cells (used from as received stock), and 0.5 µl of stock Calcein AM for live cells. After 20 min incubation with the Live-Dead cocktail, six micrographs per well were taken using a Nikon Eclipse Ti–U fluorescence microscope. The images were acquired using 495/515 nm filter for Calcein, and 528/617 nm filter for Ethidium Homodimer-1 (EthD-1). The number of live or dead cells were counted in each image to analyze cell cytotoxicity after laser irradiation at different time points.

### In vitro PA imaging

A Vevo Lazr device (VisualSonics) with LZ550 transducer were used to acquire the photoacoustic images. 20 μl of PTNP were inserted into 0.5 mm diameter polyethylene tubing which was then covered in 2–3 cm of water. PTNP were prepared at different PTNP concentrations (0, 0.06, 0.12, 0.25 and 0.5 mg ml^-1^). The transducer distance from sample was 1.1–1.5 cm.

The PA imaging of PTNP within skin biofilms was also carried out using the Vevo Lazer device and LZ550 transducer. The S. aureus bacteria were cultured in 96 wells using Bacto Tryptic Soy Broth (TSB) (BD #211825) as a growth media. After 24 h, the biofilm bacteria were incubated with different PTNP concentrations (0, 0.12, 0.25 and 0.5 mg ml^-1^) for another 24h. Then the biofilm treated with PTNP was washed 3 times and were covered in 2-3 cm of water. The ultrasound gain and PA gain were adjusted at +27 dB and 22 dB, respectively.

## Animal and infectious disease models

### Ex vivo efficacy of PTNP for oral biofilms

An established rodent model that mimics dental caries, including S. mutans infection of rat pups, was used for ex vivo experiments for oral infectious disease (90, 93). Briefly, 15-day-old female Sprague−Dawley rat pups were purchased from Harlan Laboratories. First, the animals were infected orally with actively growing (mid-logarithmic) culture of S. mutans UA159, and their infection was confirmed via oral swabbing. The animals’ diet was based on the NIH cariogenic diet 2000 (TestDiet) and 5% sucrose water ad libitum. The animals were categorized into five treatment groups. The treatment groups were: (1) control (0.1 M NaAc buffer, pH 4.5), (2) PTNP only (0.25 mg ml^-1^), (3) laser only, (4) PTNP and laser irradiation and (5) chlorhexidine. The animals’ physical appearance was monitored daily, and the body weights were recorded weekly. The experiment proceeded for 3 weeks (21 days). At the end of the experimental period, the animals were sacrificed, and their jaws were surgically removed and aseptically dissected. Jaws with infected teeth were placed in a solution of PTNP (at 0.25 mg ml^-1^), chlorhexidine or vehicle for 10 min and washed in DI water (the Jaw was dipped in DI water 3 times), which was immediately followed either by NIR laser irradiation (at 808 nm wavelength and 2 W cm^-2^) for 1 min or mock irradiation. Some jaws were used for PA imaging and others were used for CFU counting and biofilm biochemical assessment. For the latter, the plaque−biofilm samples were removed from the jaws and teeth using sonication and prepared for analysis (93).

### In vivo efficacy of PTNP in skin biofilm

To test the ability to kill bacteria in vivo, a modified S. aureus (Xen36 luminescent strain, a pathogen known to cause wound infections) delayed inoculation wound infection model was developed (84), (87). Male 8–12-week-old C57BL/6J mice were bred in the vivarium at Stanford University. The ones that underwent surgery received additional Supplical Pet Gel (Henry Schein Animal Health, Cat. No. 029908). Mice are divided into 5 groups: (1) control, (2) PTNP only, (3) laser only, (4) PTNP + laser and (5) topical antibiotic positive control (gentamicin). Mice were anesthetized by delivering 1%–3% isoflurane and weighed to obtain pre procedure weight. Mice were shaved using an electric shaver on the dorsal area, followed by applying a thin layer of hair removal lotion for 30 s to remove extra hair. Then the area was cleaned with gauze moistened in warm water. A day after hair removal, an excisional wounding procedure was performed. First, the dorsal area of the mouse was injected with buprenorphine 0.6–1 mg/kg using a 25 G needle to reduce the pain over the next 48–72 h. After disinfecting, the surgical site was wiped with betadine and sterile alcohol swab 3 times. A 6 mm diameter excisional wound was created with a punch biopsy tool (Miltek) to make an initial incision through the left and right side of dorsal epidermis. The selected area was excised on both sides of the wound area using scissors. After washing the wounds with sterile saline, the wounds were covered with a transparent film dressing and the mice were placed in clean cages.

24 h after surgery, 1-2 × 10^8^ CFU of XEN36 bioluminescent strain S. aureus were injected through the transparent film dressing into each wound. For the subsequent 48 h, mice were monitored at least daily for wound condition and signs of distress. Meanwhile, the infected wounds were monitored with BLI to visualize the infection rate in each wound. After 48 h, the treatment procedures were carried out. For the PTNP only and PTNP + laser groups, 20 µL of 0.25 mg ml^-1^ of nanoparticles were applied to an inoculated skin wound. After an incubation time of 10 min, whilst the mice were under anesthesia, the infected wound with the associated nanoparticles, and laser only group were exposed to the laser treatment (808 nm NIR laser, at 0.7 W cm^-2^) for a duration of 5 min. Other groups were treated with the same condition, but without the NIR laser irradiation. Animals were euthanized after the treatment and in vivo imaging. Then the treated and untreated wounds were extracted for bacterial plate count. The surface temperature of wounded mice was also monitored before treatment, after 1, 2, 3, 4 and 5 min of treatment using an FLIR ONE thermal imaging camera (FLIR Systems).

## In vivo imaging

### Bioluminescence imaging

BLI was performed by using a Lago X imaging system (Spectral Instruments Imaging). Mice infected with luciferase expressing XEN36 S. aureus were anesthetized with 2.5% isoflurane and bioluminescent signals were quantified by using Aura software (Spectral Instruments Imaging). The acquisition parameters were adjusted as follows; exposure time 30 s, binning low (2), f/stop 1.2, and FOV 25. The luminescence data was analyzed using Aura software.

### In vivo PA imaging

PA was performed with a VisualSonics Vevo 2100/LAZR high-frequency ultrasound/photoacoustic imaging scanner with 256 array transducers. The experiment was carried out with skin infected male 8–12-week-old C57BL/6J mice (n = 3). The procedure was performed under anesthesia with isoflurane (1.8–3%, 2 l min^−1^ O^2^) before imaging on control and PTNP treated mice (20 µl of 0.25 mg ml^-1^) for 10 min. The images were acquired using 800 nm NIR light, with ultrasound gain of +27 dB, PA gain 35 dB, and priority 95%.

### Ex vivo PA imaging

Ex vivo PA imaging was captured using a Vevo Lazr @2100 device (VisualSonics). PA imaging was performed on infected jaws (n = 3) which were exposed to PTNP (0.25 mg ml^-1^) for 10 min, and infected jaws which had no incubation with PTNP (control). The jaws were adhered to a dish with glue and were covered with 1-2 cm water. An LZ550 transducer, 800 nm NIR light, with ultrasound gain of +27 dB, and PA gain 35 dB, were used to acquire PA images. Image analyses were performed by placing ROIs on the images of control and biofilms incubated with PTNP. Then signal-to-background ratio (SBR) values were calculated from these measurements (from six images per group).

### Statistical analysis

Statistical analyses for the experimental data were performed using a mixed-model analysis of covariance (ANCOVA) and SAS 9.5 (SAS Institute). For the in vivo experiments, an analysis of outcome measures was done with transformed values of the measures to stabilize variances. The data were then subjected to analysis of variance (ANOVA) for all sets of data.

## Acknowledgements

This work was supported by funding from NIH-NIBIB grants K99EB028838 (MH), R01-DE025848 (HK) and R01-HL131557 (DPC). C.D.V. was supported by NIH grants T32 AI007502 and K08AI151089, as well as a Doris Duke Physician Scientist Fellowship. JCH was supported by a Brody Postdoctoral Fellowship. We acknowledge Biao Zuo, Susan Schultz, and Elizabeth Higbee- Dempsey from the Electron Microscopy Resource Laboratory, the Radiology Department, and the Bioengineering Department of University of Pennsylvania for their help with the TEM sample preparation, PA scanner, and PTT set-ups preparation, respectively. We also acknowledge the Stanford Center for Innovation in in vivo Imaging (SCi_3_) for using their small animal imaging facility.

## Data Availability

The data that support the findings of this study are available in the text and supplementary file. Additional data related to this paper may be requested from the authors.

## Conflicts of interest

There are no conflicts to declare.

## Supplementary information

Additional data is provided which shows TEM images, photostability of PTNP, in vitro and in vivo PA imaging, and analysis of PTT of infected wounds and healthy mice. This material is available free of charge via the Internet at http://pubs…

## Notes

### Competing Interest Statement

The authors have declared no competing interest.

