## Supplemental File for "Theranostic gold in a gold cage nanoparticle for photothermal ablation and photoacoustic imaging of skin and oral infections"

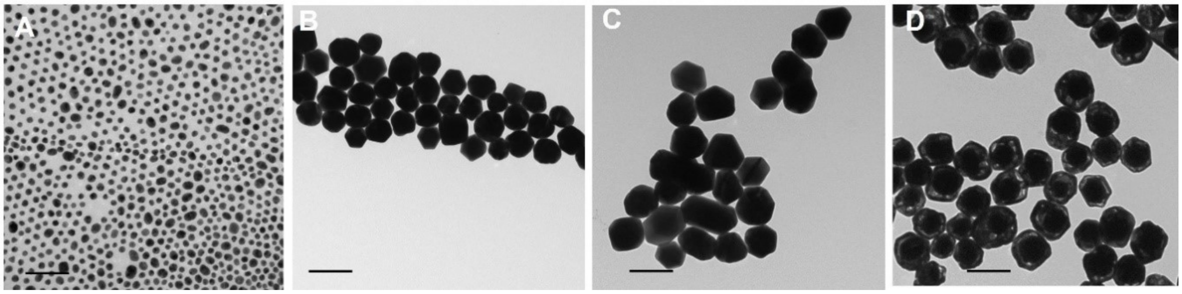


**Figure S1.** TEM of A) Au seeds, B) the 78 nm Au cores, C) Au@Ag core–shell structures, D) Au@Au core-cage structures. The scale bar is 100 nm in each panel.

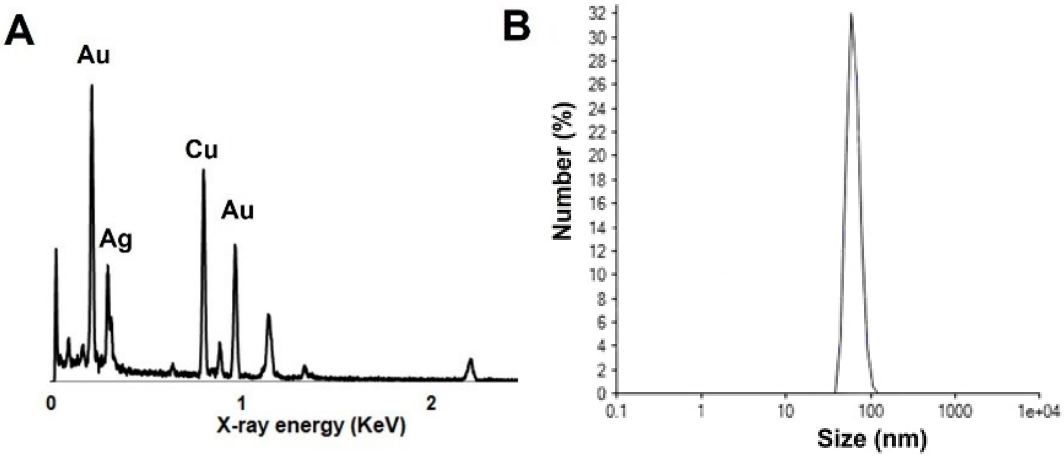


**Figure S2.** Characterization of PTNP structures. A) EDS spectra of PTNP (To measure the EDS spectra, the samples were prepared on TEM grids which are made of copper), and B) Hydrodynamic diameter distribution of PTNP structures.

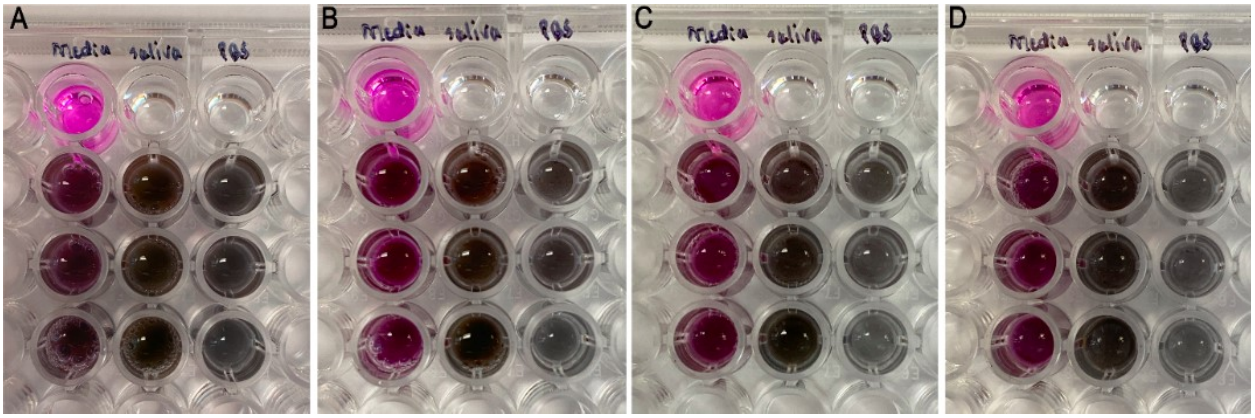


**Figure S3.** PTNP suspended in HaCaT cell media, saliva, and PBS for A) 1 day, B) 3 days, C) 8 days, and D) 15 days.

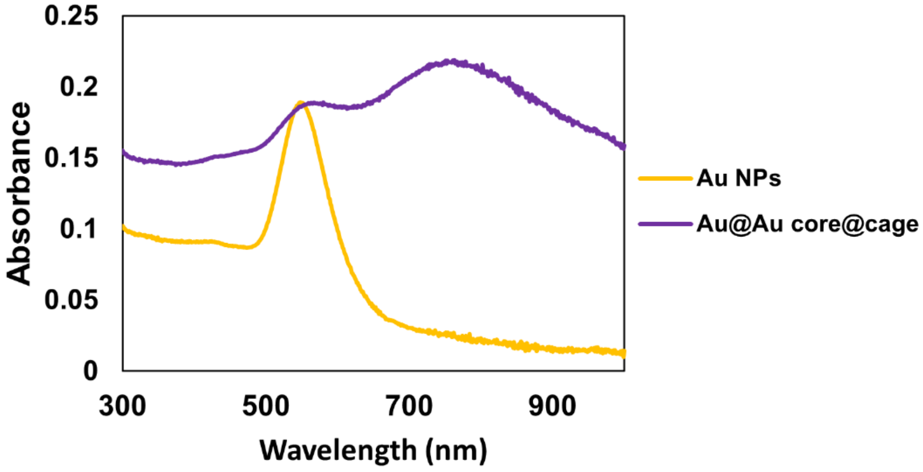


**Figure S4**. UV-vis spectrum of AuNPs and Au@Au core@cage nanoparticles (PTNP).

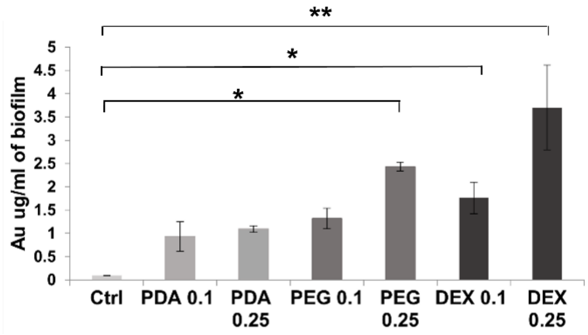


**Figure S5**. The uptake of DEX, PEG and PDA coated PTNP (in different concentration of 0, 0.1, 0.25 mg/ml) into *S. mutans* biofilms formed on sHA. *P< 0.05, **P<0.005.

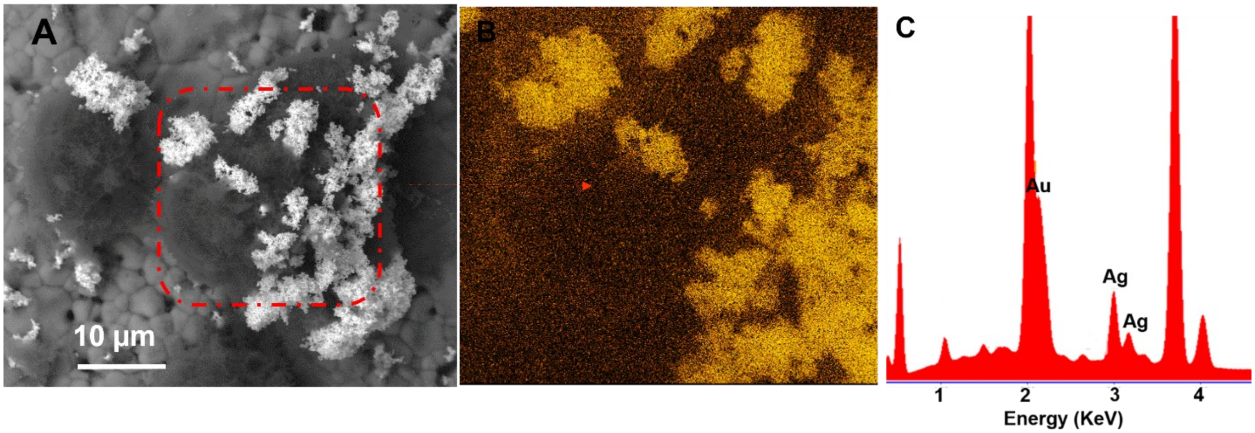


**Figure S6.** PTNP uptake by s. mutans bacteria. A) SEM in backscattered electron (BSE) mode showing the morphology of PTNP-treated biofilm. B) BSE/EDS map showing gold (yellow)

distribution on biofilms. C) Energy dispersive X-ray spectroscopy (EDS) elemental analysis of PTNP structures.

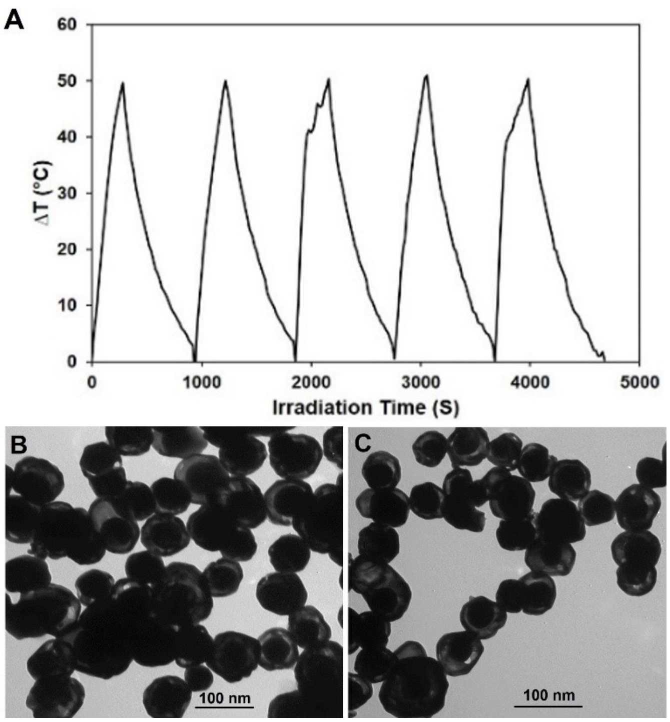


**Figure S7.** Photostability of PTNP. A) Temperature change of PTNP during five laser irradiation cycles (each irradiation time is 5 min). TEM of irradiated PTNP sample B) before, and C) after five cycle laser irradiations.

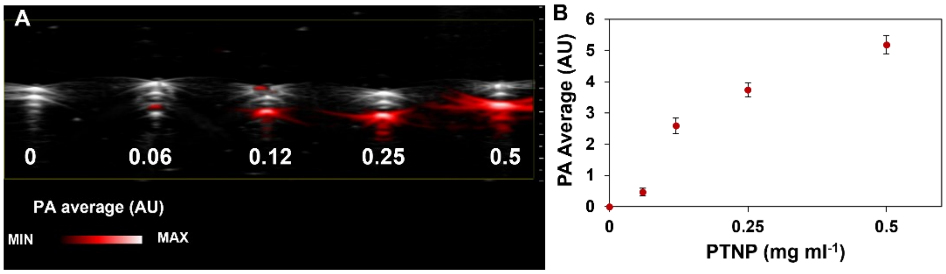


**Figure S8**. In vitro PA imaging of PTNP, A and B) PA contrast enhancement at different concentrations.

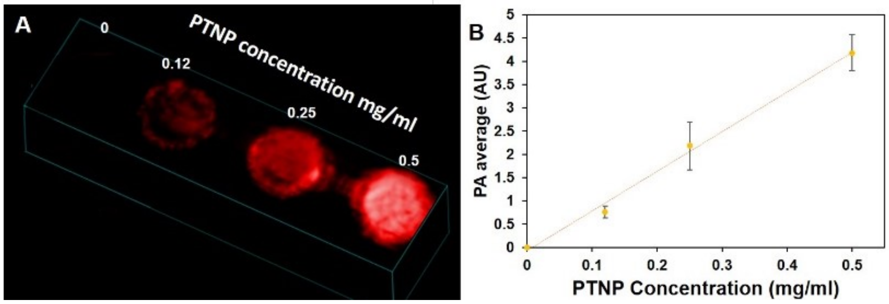


**Figure S9**. In vitro PA imaging of PTNP, A and B) Signal enhancement of PTNP incubated with S. aureus biofilm at different concentrations.

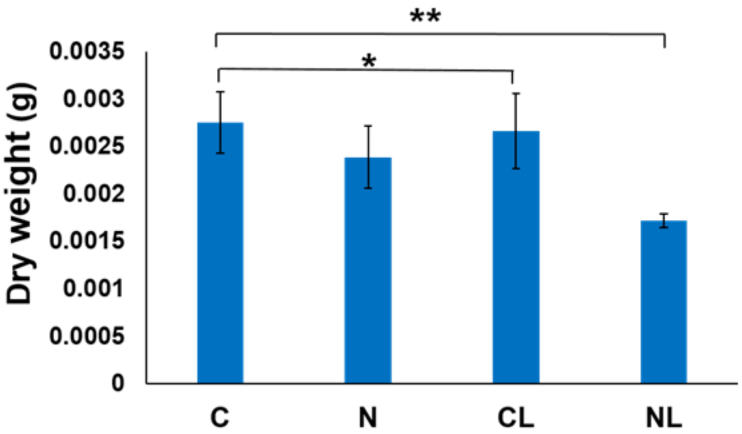


**Figure S10**. Dry weight of biofilms after the treatments. In the graphs C, N, CL, NL denote control, PTNP, control + laser, and PTNP + laser respectively. *P < 0.05, **P < 0.0001.

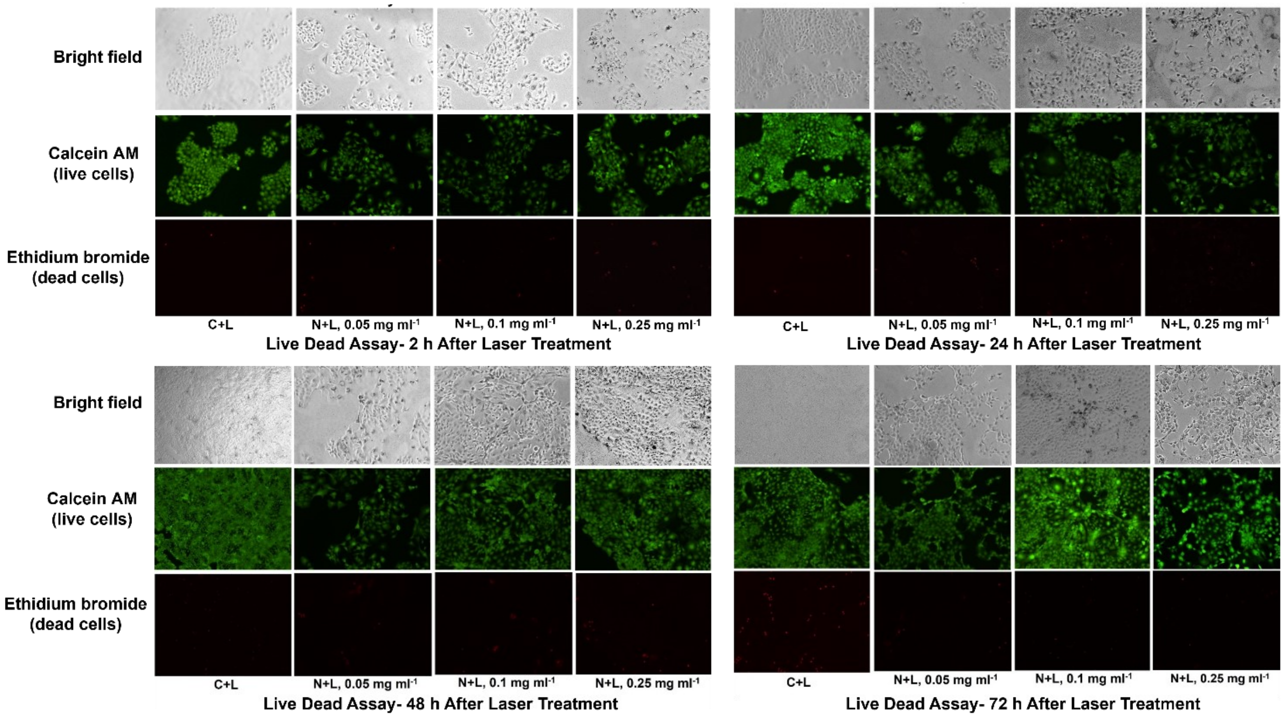


**Figure S11**. Live-Dead staining image of HaCaT cells at 0.5W laser irradiation after 2h, 24h, 48h, and 72h.

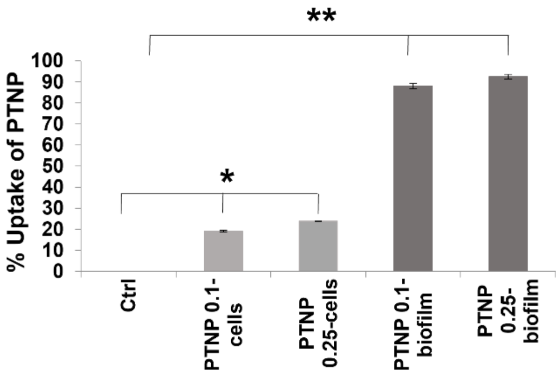


**Figure S12** The uptake of DEX coated PTNP (at concentrations of either 0.1 or 0.25 mg/ml) into S. mutans biofilms and epithelial cells. *P< 0.05, **P<0.005

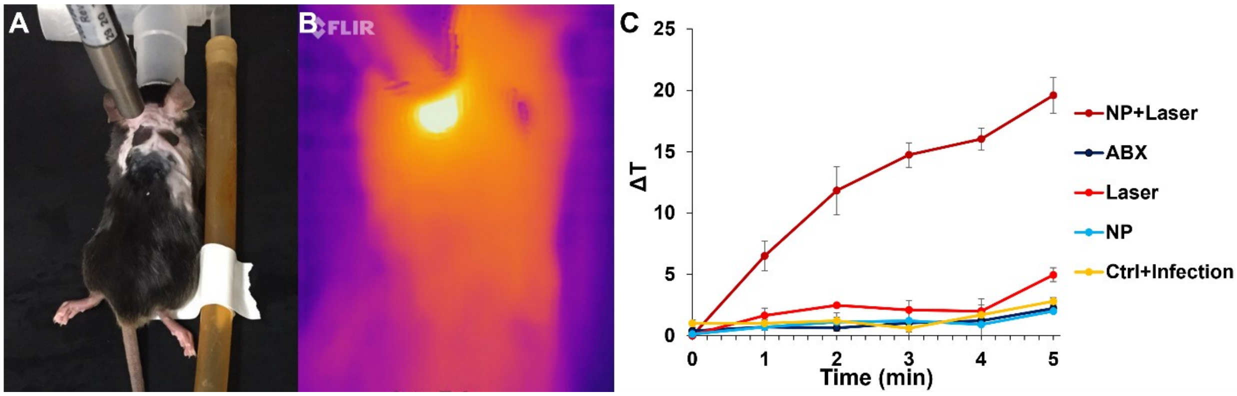


**Figure S13**. Infected wounds were administered PTNP (20 µl of 2.5 mg $ml^{-1}$, topical treatment); and ABX. 10 minutes later, infected wounds of mice also received laser irradiation (0.7 W $cm^{-2}$, 5 min). n = 4 mice per group, for a total of five groups. A) A photograph of the procedure. B) Thermographic image of an infected wound and its surroundings during the treatment. C) Image analysis of thermal imaging data during the treatment, ΔT shows the difference between wound temperature and animal body temperatures.

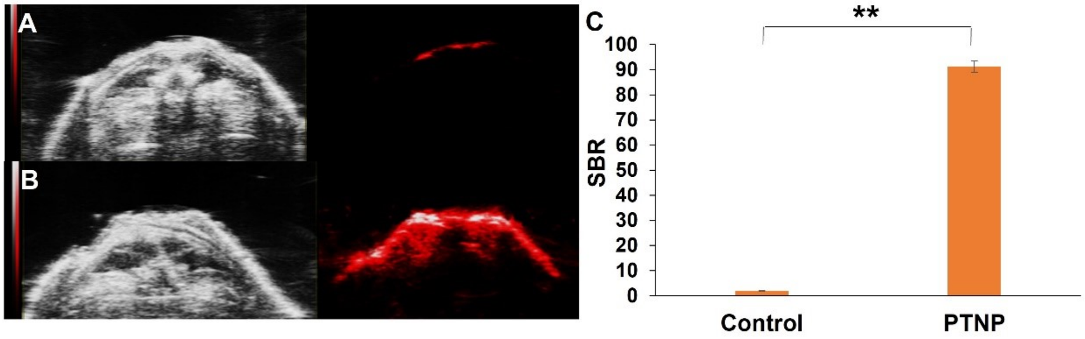


**Figure S14**. In vivo PA imaging of infected wounds. A) Control and B) PTNP treated. Left: greyscale ultrasound images, and right: PA images. C) SBR of the infected wounds with and without incubation with PTNP. ** indicate statistically significant differences at P<0001.

### Evaluation of Photothermal Conversion Efficacy

The photothermal conversion efficacy of PTNP was calculated using the following equation:

$$\eta = (hS\Delta Tmax\text{-}Qs)/\ I(1\text{-}10^{-A808}) \quad (1)$$

$$Ts = mD\ CD/hS \quad (2)$$

Where h is the heat transfer coefficient, S is the surface area of the container, Tmax is the maximum steady temperature of the solution of the PTNP (i.e., 71.2 °C), and environmental temperature (TSurr) was 18 °C. I is the laser power (2 W), A808 is the absorbance of the PTNP at 808 nm (A= 0.17), and Qs express heat associated with the light absorption by the solvent. The variable Ts is the sample-system time constant, and mD and CD are the mass (1 g) and heat capacity (4.2 $J \cdot g^{-1} \cdot °C^{-1}$) of the deionized water used as the solvent. From equations (1) and (2), the η value of the PTNP was calculated to be 77%, which is comparable with gold nanostar (78%)[1], gold nanoporous nanoshell (75.5%)[2] and higher than gold nanocages (53.6%)[3], gold nanorings (42%)[4], gold nanoshells (41.4%)[5], gold nanorod (21.3%)[6] graphene oxide (25%)[7], Cys-CuS NPs (38%)[5] Au@ Cu2–x S Core@ Shell (52%)[8].
